# *In vitro* and *in vivo* synergy of Vancomycin and β-lactams against drug-resistant *Mycobacterium tuberculosis* and Non-tuberculous mycobacteria

**DOI:** 10.1101/2025.03.24.644967

**Authors:** Alok Kumar Singh, Pradip Malik, Atri Mukhopadhyay, Mohmmad Imran, Mohammad Naiyaz Ahmad, Juned Ali, Sangita Pramanik, Reetu Maindolia, Aditi Ghosh, Pratiksha Karaulia, Umesh D Gupta, Kalyan Mitra, Sidharth Chopra, Arunava Dasgupta

## Abstract

The unabated increase in antimicrobial resistance has underlined the importance of identifying novel drug combinations which eliminate infections more potently and likely reduce the emergence of resistance. In this context, we have identified Vancomycin and many β-lactams as being potently active against drug-resistant *Mycobacterium tuberculosis* and non-tuberculous mycobacteria including *M. abscessus,* emerging as pathogens of concern owing to their inherent drug resistance profile. In this study, we have identified combinations of Vancomycin, a glycopeptide and β-lactams, especially Ceftriaxone, Ceftazidime and Meropenem, in the presence or absence of Sulbactam, a β- lactamase inhibitor, as possessing potent antimicrobial activity against several drug-resistant mycobacterial strains. The combination of Vancomycin and β-lactams exhibited potent bactericidal activity and reduced the bacterial load better than either drug alone. The molecular basis of synergy was mediated by increase in permeability of mycobacterial cell as demonstrated by ethidium bromide assay. In the murine model of mycobacterial infection, synergistic combination of Vancomycin and β-lactams outperformed clinically utilized drugs including Isoniazid, Rifampicin and Ethambutol against *M. tuberculosis* and Amikacin, Clarithromycin against *M. abscessus*. The combinations caused a significant reduction in bacterial load in various organs in *M. tuberculosis* and *M. abscessus* infected mice. Thus, the synergistic combination of Vancomycin and β-lactams could potentially be utilized for treatment of recalcitrant mycobacterial infection especially those caused due to drug-resistant pathogens.

---

Tuberculosis (TB) caused by *Mycobacterium tuberculosis* (Mtb) has returned back to leading cause of death worldwide due to single infectious agents with more than ∼1.25 million deaths and ∼8.2 million new cases reported in 2023 (1). The multi drug-resistant (MDR) or Rifampicin-resistant (RR) TB still remains a critical cause of health concern as only 44% of total 4000,000 people estimated to have developed MDR/RR TB were diagnosed and treated. Although the treatment success rate for drug-resistant TB has increased from 63% in 2022 to 68% in 2023 but still significantly below the target of the “End TB strategy” of WHO (1). In addition to Mtb, the genus *Mycobacterium* comprises of more than 170 species commonly known as non-tuberculous mycobacteria (NTMs). Among them, some including *Mycobacterium abscessus*, are rapidly emerging as a threat to healthcare due to its notorious drug resistance profile and significant increase in number of reported cases worldwide particularly in immuno-compromised individuals as well as patients with cystic fibrosis (2–5). Like Mtb, these pathogens also cause pulmonary and extrapulmonary infections. (6–7)

The treatment of various drug resistant mycobacterial infections, including TB, has always been challenging due to multiple reasons including multiple drugs and long periods of chemotherapy amongst others but with the emergence of antimicrobial resistance (AMR), the situation has turned alarming with rapid decrease in patient compliance, thus stressing global healthcare systems (8,9). This chemotherapeutic situation is much worse for NTM infection especially for *M. abscessus* pulmonary infection as they are intrinsically resistant to most frontline antimicrobials and inducibly resistant to others (10). In addition, infections caused due to NTMs including *M. abscessus* are often misdiagnosed as TB resulting in usage of gratuitous and non-effective antibiotics for more than a year not only results in toxic side effects but also contributing to AMR. To overcome the situation, there is an urgent need for the discovery and development of new therapeutic interventions which would enable escape from the existing AMR mechanisms. This urgency is reflected in a robust TB drug discovery pipeline but there is almost negligible NTM-specific discovery pipeline (11). Upon closer examination, the TB discovery pipeline is replete with “me too” drugs for which sizeable resistance mechanisms already exist, thus jeopardizing the eventual containment of this highly infectious disease (12).

The bacterial peptidoglycan (PG) biosynthesis is considered to be a highly conserved, specific, antibacterial target in mycobacteria as well as various Gram positive and Gram negative bacterial pathogens, a large number of clinically utilized drugs targeting its various components (13,14). Despite their wide utilization and well-studied pharmacokinetic and pharmacodynamic properties, historically, the majority of PG inhibitors such as β-lactams, carbapenems and glycopeptides have not been clinically utilized against mycobacterial infections because of the inherent presence of β-lactamase. However, in recent times β-lactams (BL) in combination with β-lactamase inhibitor (BLI) have shown activity against clinical isolates of MDR and XDR Mtb as well as *M. abscessus* (15–20). It is noteworthy to mention that during the preparation of this manuscript, two publications reported *in vitro* combinatorial activity of BLs and BLI against Mtb and *M. abscessus* (21,22). Moreover, recently it has been shown that carbapenems as well as ceftriaxone (CRO) and cefotaxime demonstrated promising binding affinity towards all Penicillin-Binding Proteins (PBPs) while ceftazidime (CAZ) and cefoxitin moderately bind to PBPs (23). In addition, it was also demonstrated that the effect of the combination of sulbactam (SUL) and ampicillin was better than clavulanate/amoxicillin and tazobactam/pieracillin against six pathogenic mycobacteria (24). We hypothesized that a combination of glycopeptide and BL may synergistically weaken the cell wall of the pathogen, leading to potent efficacy. In this line, we present evidence in this manuscript that the combination of vancomycin (VAN) and various BLs, especially high-PBP binders meropenem such as (MEM) and CRO and a moderate-PBP binder, CAZ, in the presence or absence of SUL, a BLI do synergize *in vitro* and *in vivo* against drug-resistant mycobacterial pathogens and could constitute a potential new therapeutic option. The addition of a BLI such as Sulbactam did not significantly increase the synergistic activity of the combination *in vivo*, suggesting that the combination of glycopeptide and a PBP binder overcome the intrinsic resistance present in mycobacterial pathogens.

## Results

### MIC of vancomycin and β-lactams against different mycobacterial pathogens

The anti-mycobacterial potency of VAN and different BLs were determined against a panel of mycobacterial strains utilizing broth microdilution and the MICs are listed in Table 1. The panel consisted of Mtb H37Rv ATCC 27294, INH resistant Mtb ATCC 35822, RIF resistant Mtb ATCC 35838, STR resistant Mtb ATCC 35820, EMB resistant Mtb ATCC 35837, *M. chelonae* ATCC 35752 and *M. abscessus* ATCC 19977, *M. abscessus* MC1815, *M. abscessus* DJO 4274; representing a mix of drug-susceptible and drug-resistant mycobacterial pathogens. As can be seen in Table 1, all mycobacterial strains were susceptible to VAN (MIC 1-4 µg/ml) whereas the MIC increased to 16-32 µg/ml for *M. abscessus* strains. On the other hand, all Mtb strains, including the drug-resistant ones, were more susceptible to CRO, CAZ, SUL and MEM (MIC 2-32 µg/ml) than NTMs (MIC 4-512 µg/ml). This high MIC for NTMs is consistent with earlier reports citing the variable efficacy of β-lactams against various mycobacteria (25). It is noteworthy to mention that the clinical strains of *M. abscessus, M. abscessus* MC1815 and DJO4274 were found to resistant to several antibiotics including Amikacin and Clarithromycin (Table 1).

**Table 1:** Minimum inhibitory concentration (MIC) of various drugs against mycobacterial pathogens in µg/ml.

| Bacterial strain |  | MIC (µg/ml) |  |  |  |  |  |  |  |  |  |  |  |  |  |  |  |  |  |  |  |
| --- | --- | --- | --- | --- | --- | --- | --- | --- | --- | --- | --- | --- | --- | --- | --- | --- | --- | --- | --- | --- | --- |
|  |  | VAN | CRO | CAZ | MEM | SUL | IPM | ETP | PTZ | DRP | INH | RIF | EMB | STR | BDQ | CFZ | MXF | LZD | LVX | AMK | CLR |
| Mtb | H37Rv ATCC 27294 | 4 | 4 | 32 | 4 | 16 | NT | 4 | >64/32 | 4 | 0.03 | 0.03 | 0.5 | 0.125 | 0.06 | 0.125 | 0.125 | 0.125 | 0.125 | 0.125 | 2 |
|  | RIF<sup>Res</sup> ATCC 35838 | 4 | 2 | 32 | 2 | 16 | NT | 8 | >64/32 | 8 | 0.03 | 64 | 1 | 0.125 | 0.06 | 0.125 | 0.125 | 0.25 | 0.125 | 0.125 | NT |
|  | INH<sup>Res</sup> ATCC 35822 | 4 | 2 | 32 | 2 | 16 | NT | 4 | >64/32 | 4 | 64 | 0.03 | 0.5 | 0.25 | 0.06 | 0.125 | 0.125 | 0.125 | 0.25 | 0.25 | NT |
|  | EMB<sup>Res</sup> ATCC 35837 | 1 | 2 | 32 | 2 | 16 | NT | 4 | >64/32 | 8 | 0.03 | 0.03 | >64 | 0.25 | 0.06 | 0.125 | 0.125 | 0.125 | 0.125 | 0.125 | NT |
|  | STR<sup>Res</sup> ATCC 35820 | 1 | 2 | 32 | 2 | 16 | NT | 8 | >64/32 | 8 | 0.03 | 0.03 | 2 | >64 | 0.06 | 0.125 | 0.125 | 0.125 | 0.25 | 0.25 | NT |
| NTM | *M. chelonae* ATCC 35752 | 2 | 256 | 512 | 4 | 256 | NT | NT | NT | NT | NT | NT | NT | NT | NT | NT | NT | NT | 0.06 | 0.5 | 1 |
|  | *M. abscessus* ATCC 19977 | 16 | 256 | 512 | 16 | 256 | 8 | 8 | 128/32 | 16 | >64 | 32 | 32 | >64 | 0.125 | 1 | 0.25 | 4 | 2 | 8 | 2 |
|  | *M. abscessus* MC1815 | 32 | 32 | 256 | 16 | 256 | NT | 128 | >512/256 | 16 | >64 | 64 | 64 | >64 | 0.25 | 2 | 8 | NT | 64 | 64 | >512 |
|  | *M. abscessus* DJ 44274 | 32 | 16 | 512 | 8 | 128 | NT | 64 | 128/64 | 8 | >64 | >64 | 64 | >64 | 0.5 | 4 | 32 | NT | 32 | >1024 | 512 |

**Abbreviation:** Vancomycin (**VAN**), Ceftriaxone (**CRO**), Ceftazidime (**CAZ**), Meropenem (**MEM**), Imipenem (**IPM**), Sulbactam (**SUL**), Isoniazid (**INH**), Rifampicin (**RIF**), Ethambutol (**EMB**), Streptomycin (**STR**), Bedaquiline (**BDQ**), Clofazimine (**CFZ**), Moxifloxacin (**MXF**), Linezolid (**LZD**), Levofloxacin (**LVX**), Amikacin (**AMK**), Clarithromycin (**CLR**), Ertapenem (**ETP**), piperacillin-tazobactam (**PTZ**), Doripenem (**DRP**), Not Tested (**NT**).

### Vancomycin synergizes with β-lactams against mycobacterial pathogens *in vitro*

To determine whether VAN and BLs synergize against various mycobacterial pathogens, Fractional inhibitor concentration (∑FIC) was determined utilizing the classical checkerboard method and results are tabulated in Table 2. As can be seen in Table 2, VAN strongly synergized with MEM, CRO and CAZ against all mycobacterial strains tested irrespective of their drug-resistance profile (∑FIC ≤ 0.5).

**Table 2:** Fractional inhibitory concentration of Vancomycin and β-lactams against mycobacterial pathogens.

| Bacterial strains | Combination of Van with BLs (MIC in μg/ml) | | | | | |
| --- | --- | --- | --- | --- | --- | --- |
|  | VAN + MEM |  | VAN + CRO |  | VAN + CAZ |  |
| | ΣFIC | Inference | ΣFIC | Inference | ΣFIC | Inference |
| Mtb H37Rv ATCC 27294 | 0.09375 | Synergy | 0.5 | Synergy | 0.5 | Synergy |
| RIF<sup>resistant</sup> ATCC 35838 | 0.375 | Synergy | 0.375 | Synergy | 0.375 | Synergy |
| INH<sup>resistant</sup> ATCC 35822 | 0.375 | Synergy | 0.09375 | Synergy | 0.0625 | Synergy |
| EMB<sup>resistant</sup> ATCC 35837 | 0.5 | Synergy | 0.5 | Synergy | 0.375 | Synergy |
| STR<sup>resistant</sup> ATCC 35820 | 0.5 | Synergy | 0.5 | Synergy | 0.375 | Synergy |
| *M.abscessus* ATCC 19977 | 0.375 | Synergy | 0.5 | Synergy | 0.375 | Synergy |
| *M.chelonae* ATCC 35752 | 0.5 | Synergy | 0.5 | Synergy | 0.5 | Synergy |
| *M.abscessus* MC1518 | 0.15625 | Synergy | 0.09375 | synergy | 0.14063 | Synergy |
| *M.abscessus* DJO-44274 | 0.15625 | Synergy | 0.15625 | synergy | 0.15625 | Synergy |

This observed synergy was further confirmed by combinatorial time-kill analysis both in Mtb (Fig.1) and *M. abscessus* (Fig. 2). As shown in Fig 1. A (I – III), the combinatorial effect of VAN and all the BLs tested reduced ∼7.5 log_10_ CFU in 7days compared to untreated Mtb H37Rv demonstrating equipotent efficacy to any of the front-line drugs, INH and RIF whereas the combination significantly outperformed the other first-line drug, EMB. More interestingly, in presence of SUL, a BLI the combination outperformed even INH and RIF where the CFU were eliminated below the detection level within 5-6 days (∼ 8.5 log_10_ reductions in CFU compared to untreated).

**Figure 1:**
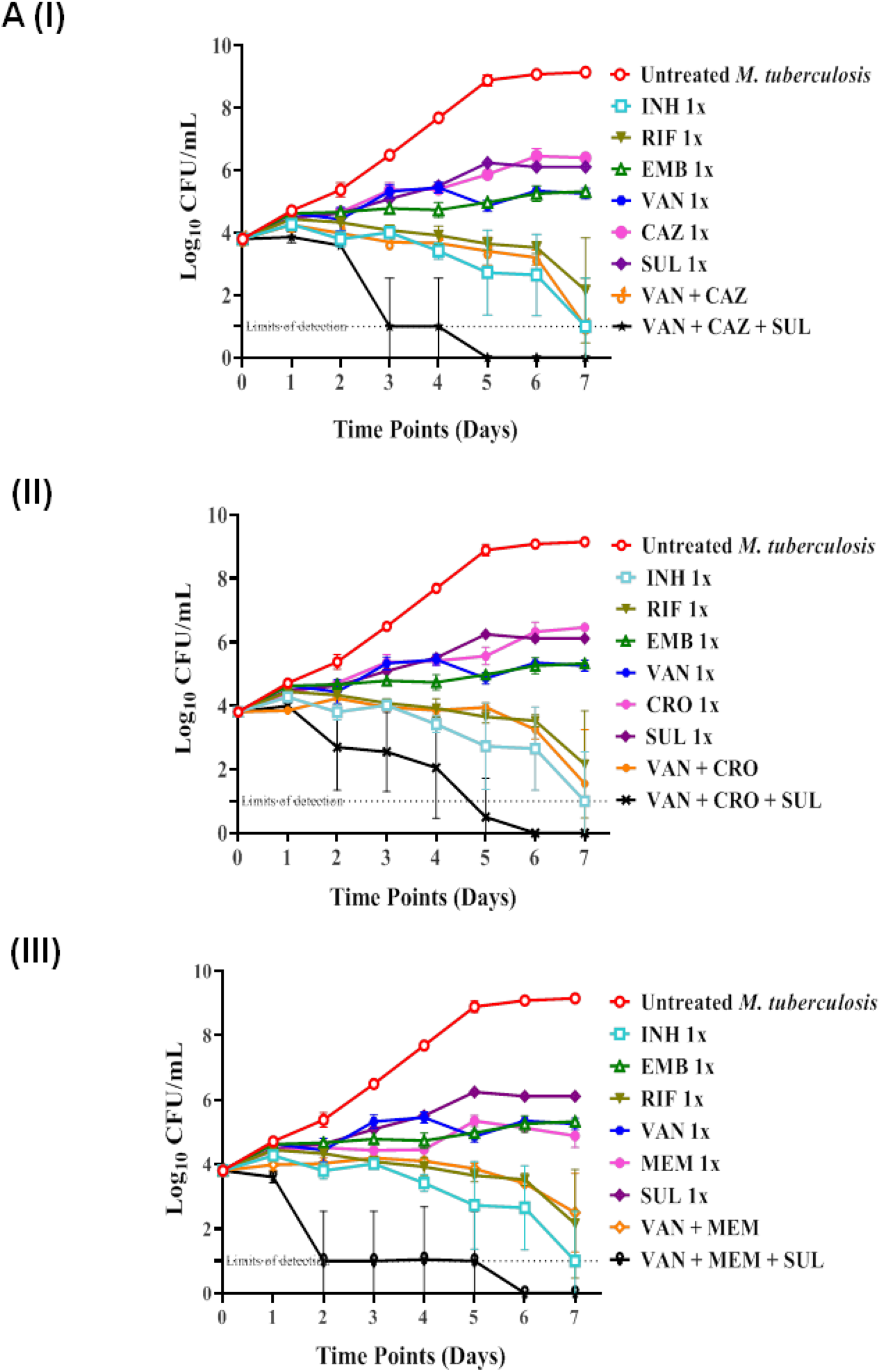
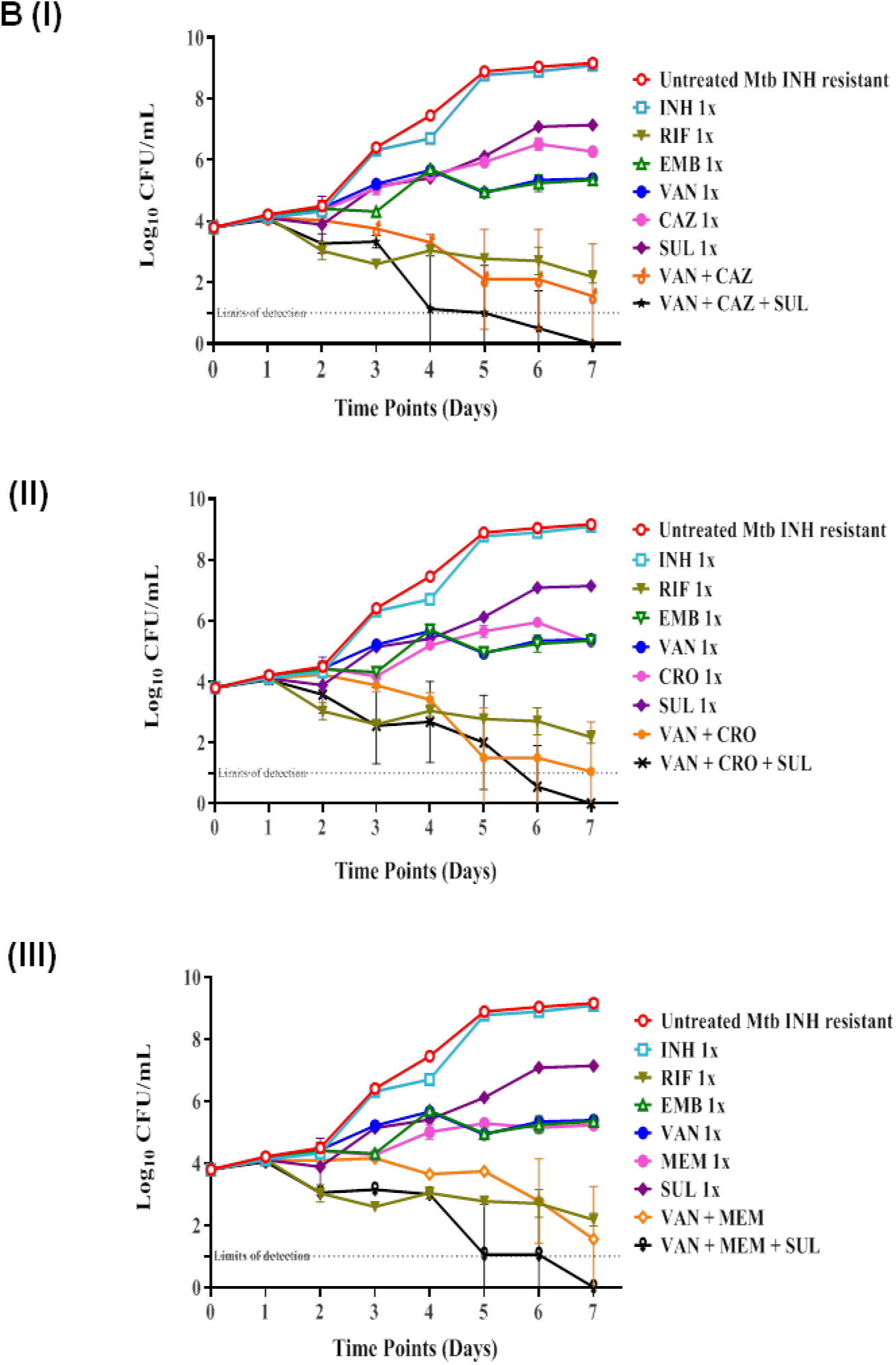
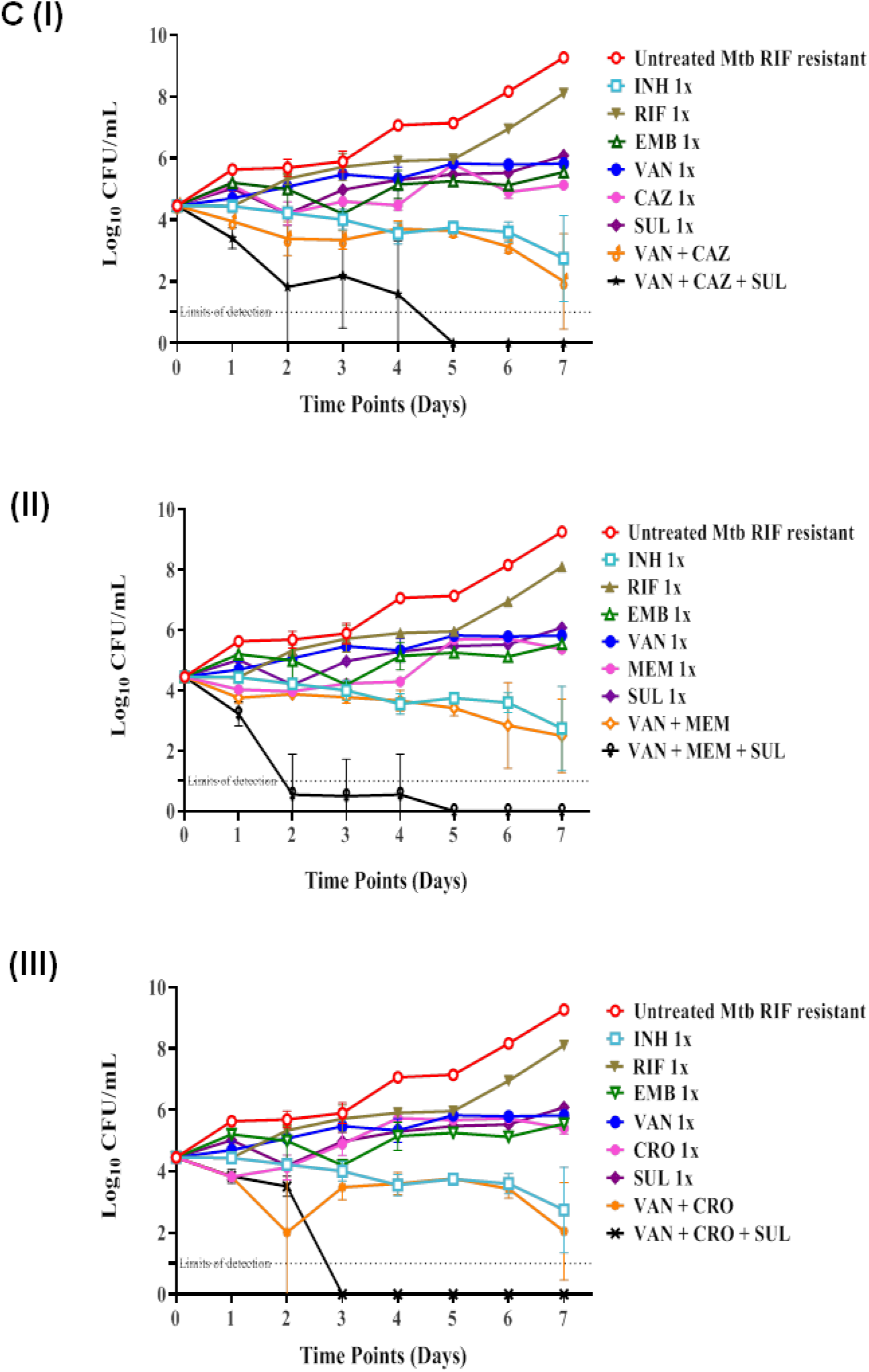
Time-kill kinetics of Vancomycin combination with beta-lactams antibiotics against various Mtb strains, (A) *M. tuberculosis* H37Rv ATCC 27294; (B) INH resistant Mtb ATCC 35822; (C) RIF resistant Mtb ATCC 35838 was done in the same set of experiments and the results were shown in three separate panels for clarity. The log_10_ CFU/ml values of control Untreated, INH, RIF, EMB, VAN, CAZ, CRO, MEM and SUL (1x MIC) are same for respective strain such as (A) *Mycobacterium tuberculosis* H37Rv ATCC 27294, (B) INH resistant Mtb ATCC 35822 and RIF resistant Mtb ATCC 35838. The error bars in all the panels represent the standard deviation of mean values derived from duplicate samples from two different set of experiments.

**Figure 2:**
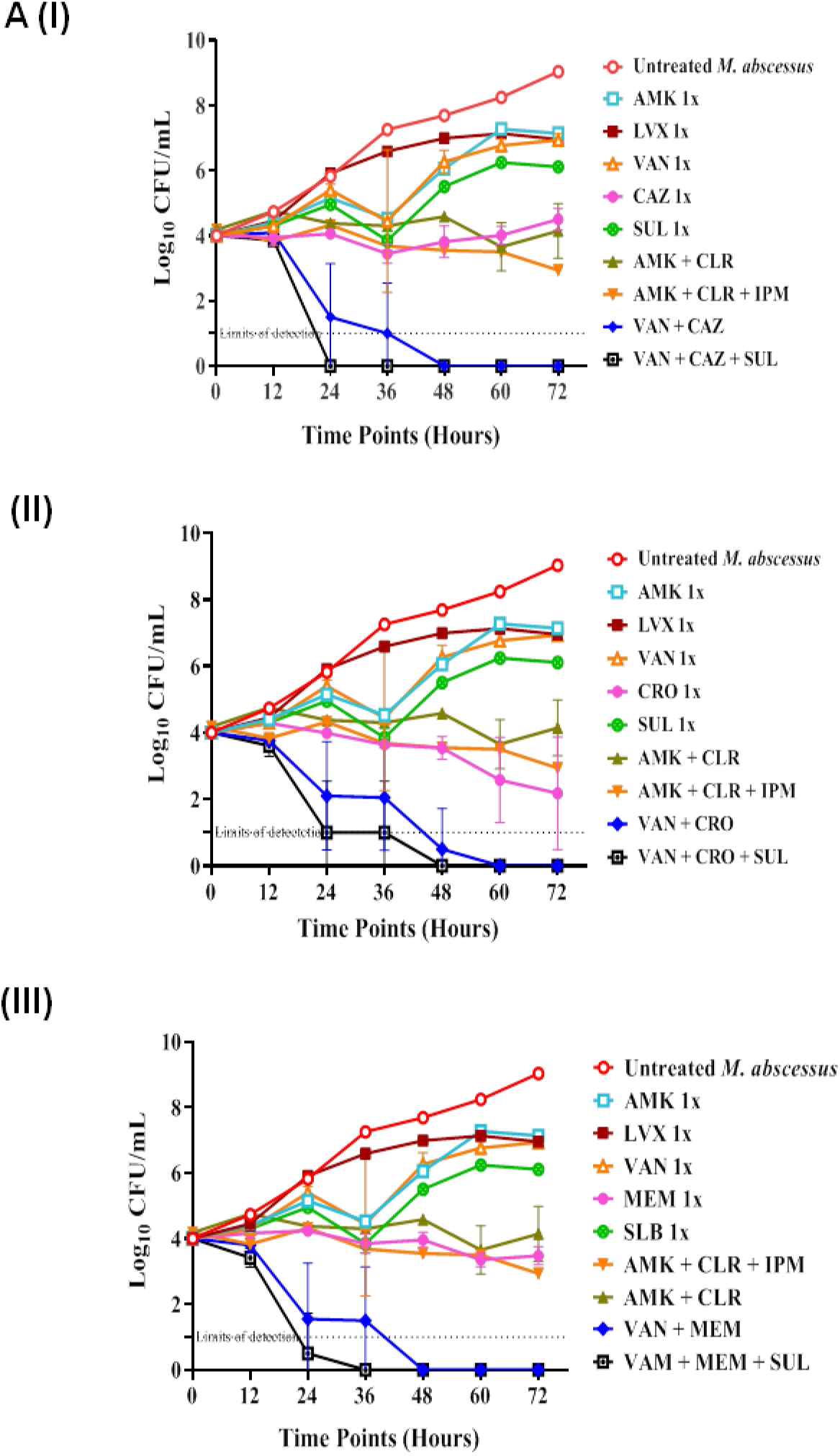
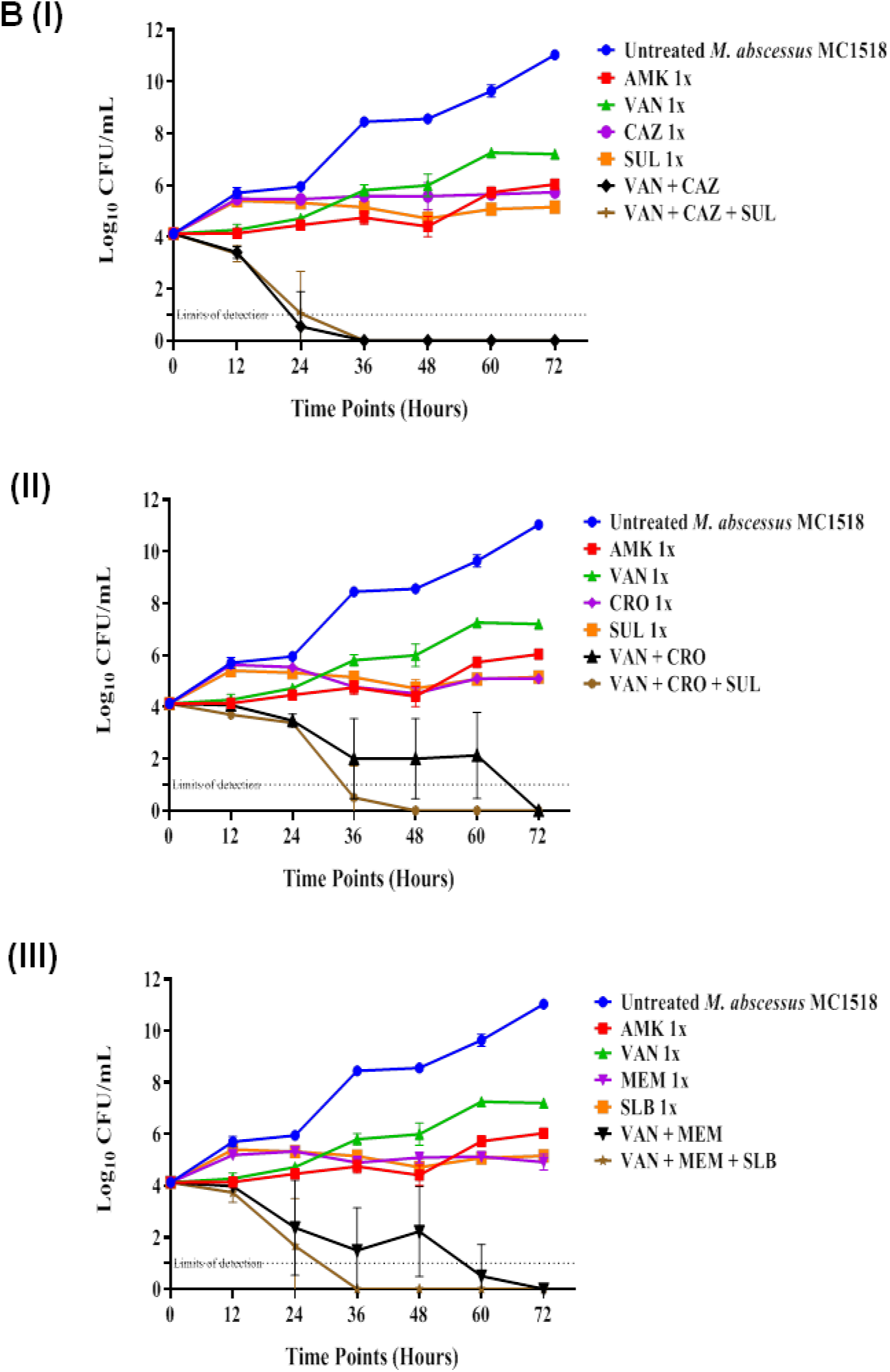
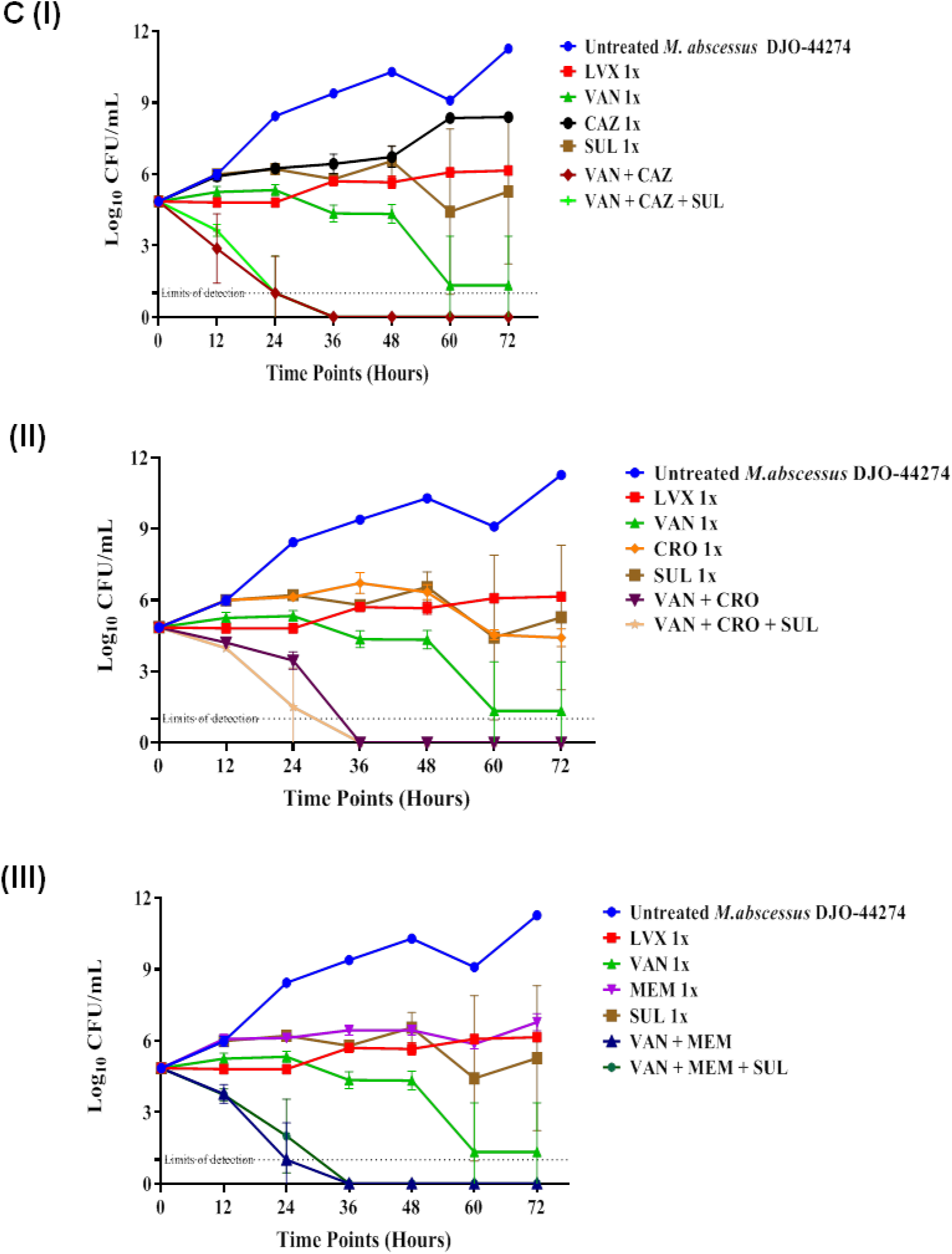
Combinations Time-kill kinetics of Vancomycin with β-lactams against various *M. abscessus* strains, (A) *M. abscessus* ATCC 19977; (B) & (C) clinical strain *M.abscessus* MC1518; *M.abscessus* DJO-44274 respectively were done in single set of experiment. Control Untreated, AMK, LVX, VAN, CAZ, CRO, MEM, SUL, AMK + CLR and AMK + CLR + IPM (1x MIC) have same log_10_ CFU/ml values for their respective pannel. The error bars in all the panels represent the SD (standard deviation) of mean values derived from replicate samples from two different set of experiments.

We further evaluated these outstanding synergistic effects of the tested combinations of VAN and BLs in presence or absence of a BLI against isogenic INH-resistant (Fig. 1B I-III) and RIF-resistant (Fig. 1C I-III) Mtb strains where similar activity were observed. The combination of VAN and BLs ± SUL comprehensively outperformed front-line anti-TB drugs against first-line drug resistant Mtb strains. The presence of a BLI enhances the activity of the combinations even further suggesting that the identified combinations of VAN, BLs and BLI could potentially be utilized against MDR TB.

The combinatorial effects were tested against extremely tough-to-treat mycobacterial strains, *M. abscessus* including drug-resistant clinical isolates and the results are shown in the Fig. 2 (A-C). First, we tested the effect of the identified combinations against *M. abscessus* ATCC19977 (Fig. 2A I-III). All the tested combinations of VAN, BLs and SUL again demonstrated excellent efficacy and comprehensively outperformed the activity of either AMK or LVX or combination of AMK + CLR and AMK + CLR + IPM which is the recommended treatment regimen for *M. abscessus* infections. The combination eliminated bacterial CFU below the detection level with ∼ 9.5 log_10_ reductions in CFU within 24-48 h.

To further assess the synergistic effect of the combinations against extremely drug resistant *M. abscessus* clinical isolates MC1518 and DJO 4274 which are resistant to almost all the antibiotics (as shown in Table 1), time-kill experiments of the identified combinations were performed against these strains (Fig. 2B I-III and 2C I-III, respectively). Remarkably, even against these notoriously resistant MC1518 and DJO 4274 strains, the combinations are highly efficacious and reduced the bacterial load ∼ 10 log_10_ CFU between 36-72 hours clearly demonstrating the potential of the combinations against several drug resistant mycobacterial infections. We noted that theaddition of SUL, a BLI significantly enhanced the efficacy of VAN +BLs. Moreover, no regrowth was observed in any of the tested combinations further demonstrating the advantages of the triple drug combinations. Collectively, these studies indicated that the identified combinations have significantly high potential as alternative therapy to macrolides, especially in those cases where inducible resistance to macrolides impairs treatment (26, 27).

### Fluorescence Microscopy bacterial viability assay against *M. abscessus* ATCC 19977

As both VAN and BLs act on the cell wall, the functional integrity of mycobacterial cell wall was assessed using BacLight live/dead assay. Viable cells are impermeable to PI but permeable to SYTO9 resulting in green florescence, whereas dead cells are permeable to PI but impermeable to SYTO9, giving red florescence. Mid-log *M. abscessus* ATCC 19977 cells were treated with 1x MIC of VAN, MEM, SUL alone and combinations of VAN+MEM and VAN+MEM+SUL for 48 h and stained with LIVE/DEAD BacLight stain, incubated in dark conditions for 15 minutes and visualized on an EVOS FLc fluorescence microscope using a 100X oil immersion objective. As seen in Fig. 3 *M. abscessus* cells treated with VAN+MEM and VAN+MEM+SUL were all dead (red) in both the cases compared to other treatments where some populations were live (green). This result indicated that VAN combinations with BLs and BLI disintegrate mycobacterial cell walls and increase permeability.

**Figure 3:**
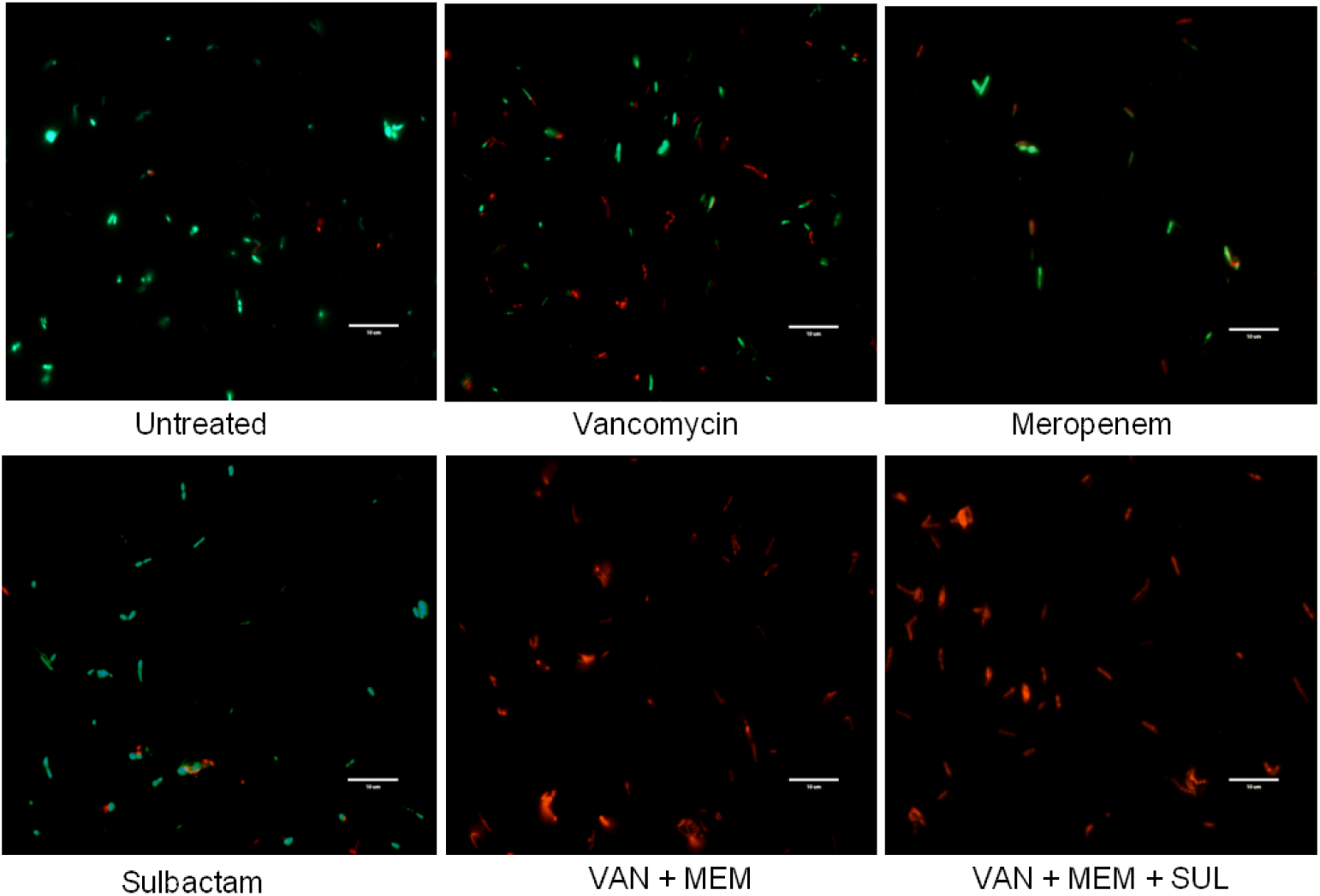
Fluorescence images of Vancomycin and β-lactams treated cells with 1x MIC, in combinations and drugs alone. After 48 hours of incubation, the untreated and treated cells are stained with PI and SYTO9. Viable cells were stained green with SYTO9 whereas the dead cells were stained red with PI. The treatment of VAN in combinations with BL (MEM) and VAN + MEM + SUL showed more dead cells (red) in comparisons to any of single drug treatment (VAN, MEM & SUL).

### Combination of Vancomycin and β-lactams increases the permeability of the mycobacterial cell wall

Since the combination of VAN and BLs exhibited high bactericidal activity, one of the hypotheses was that the combination might increase the permeability of the mycobacterial cell. Evidence to test this hypothesis was generated by studying the accumulation of EtBr in the presence of drugs alone and in combination against *M. abscessus* ATCC 19977 and results are plotted in Fig 4. A-C. As can be seen, in all cases, there is a significantly consistent increase in accumulation of EtBr in VAN + CRO, VAN + CAZ and VAN + MEM treated *M. abscessus* ATCC 19977 as compared to either drug alone. This trend is visible within 30 minutes of treatment in all cases and continues till end, thus supporting the hypothesis of increased permeability of mycobacterial cells when treated with these combinations.

**Figure 4:**
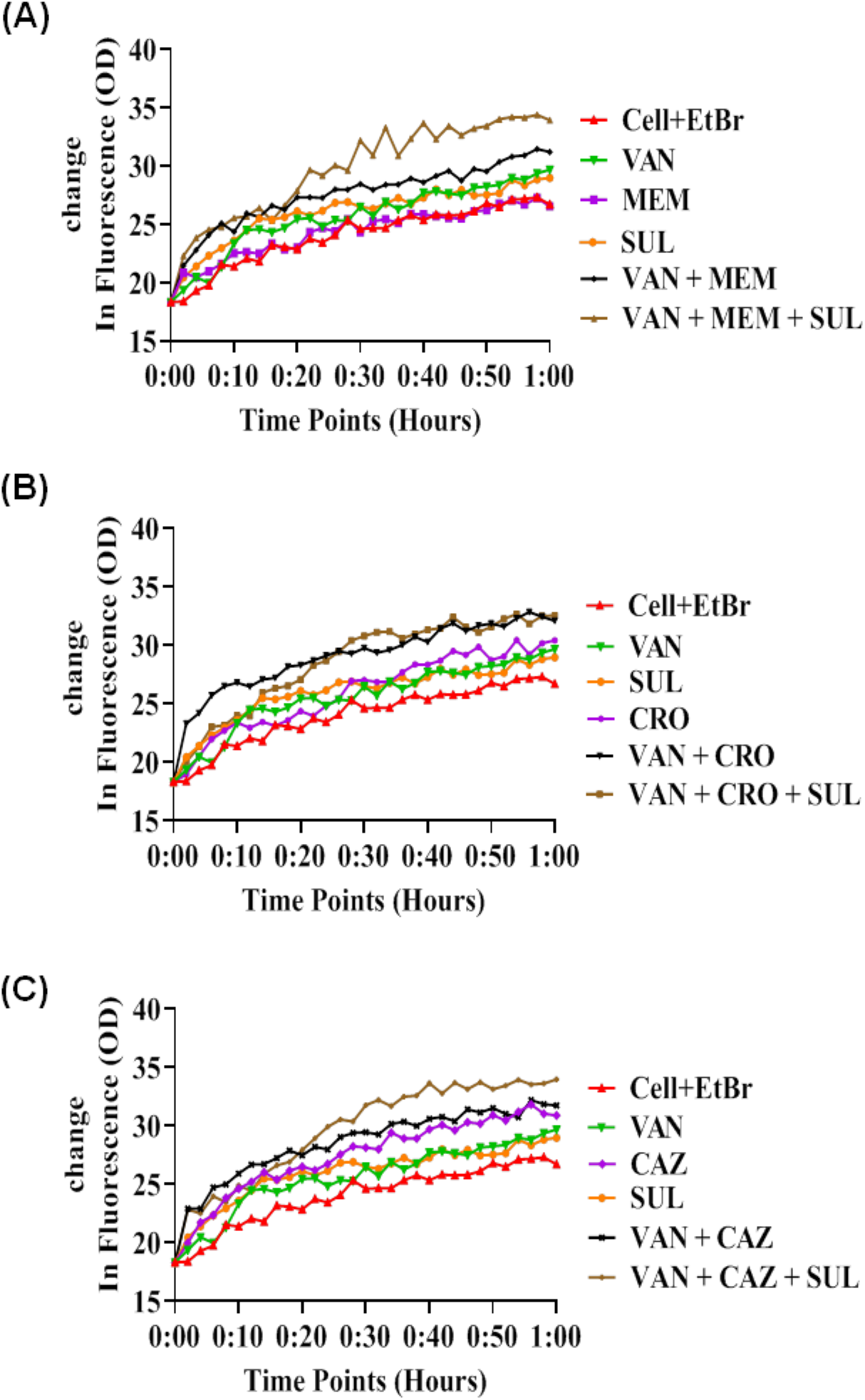
A-C. The effect of combination on cell permeability was performed against *M. abscessus* ATCC 19977, compared with drug alone. All drugs alone and in combinations were used at 0.5X MIC to not compromise cellular viability. EtBr was added to culture (O.D - OD_600nm_ of 0.4-0.6) at 10μM concentration incubated for 1 hour, relative fluorescence units (RFUs) were observed every two minutes by Excitation (530 nm)/ Emission (590 nm) filter using a multi-mode plate reader. The experiment was performed in a single set for triplicate and the mean value was plotted.

### Combination of vancomycin with β-lactams disrupts *M. abscessus* ATCC 19977 cell envelope

Scanning electron microscopy were performed to visualize the morphological and topological effects of VAN, MEM, SUL and the combinations of VAN+MEM and VAN+MEM+SUL on *M. abscessus* **(Fig.5)**. It was observed that VAN and SUL induce less visible morphological damage and the bacterial morphology is similar to untreated cells whereas MEM induces more visible morphological changes. The topological alterations and bacterial damage were more severe in cells treated with VAN+ MEM and VAN+MEM+SUL confirming that combination of VAN+BLs in presence or absence of BLI severely disrupt mycobacterial cell wall (Fig. 5).

**Figure 5:**
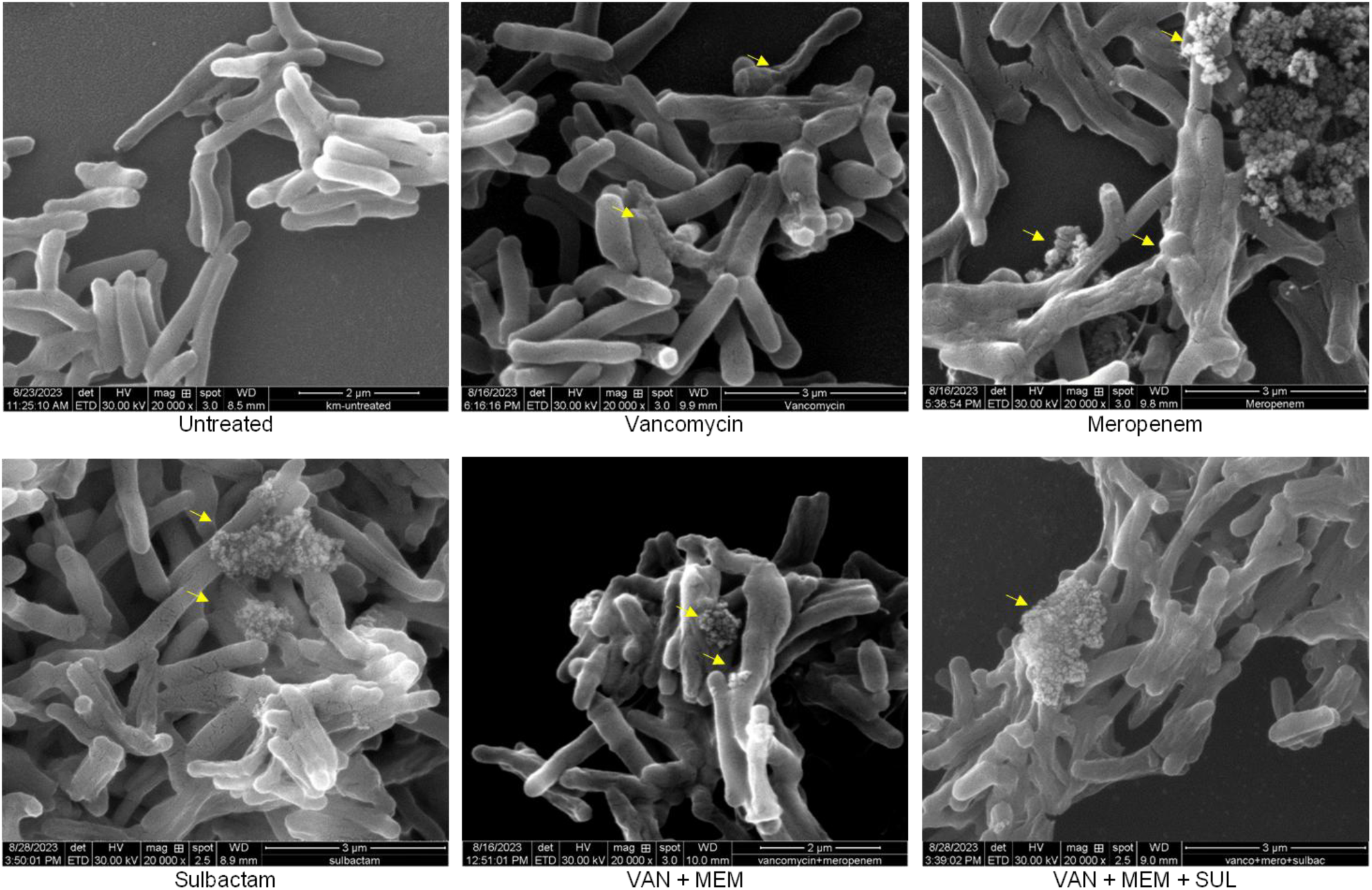
The illustration of Scanning electron microscopy images indicating the effects of drugs at 5x MIC for 8 h of treatment (drug alone and in combinations) against *M. abscessus* ATCC 19977. The effect of VAN, MEM and SUL alone does not induce visible disruptions on cell-enveloped and is similar to untreated cells but in VAN + MEM and VAN + MEM + SUL groups are showing significant alteration in topology and damages in the cell envelope.

### Intracellular killing assay against *M. abscessus* ATCC 19977

The efficacy of synergistic combinations of VAN and BLs to kill intracellular *M. abscessus* was assessed using J774A.1 cells infected with a MOI of 1:5. The infected macrophages were treated with 1x MICs of VAN, CRO, CAZ, MEM, SUL and AMK alone and the identified combinations of VAN + BLs and VAN + BLs + SUL where AMK was used as a positive control. The treated cells were then lysed and plated on ADC-7H11 agar to enumerate CFU (Fig. 6 A-C). The result demonstrated that the treatment with combination of VAN and BLs significantly reduced of bacterial load ranging from ∼ 2 log_10_ (in case of VAN + MEM) to ∼ 3.5 log_10_ (in case of VAN + CAZ) in CFU as compared to untreated control. Addition of SUL led to further decrease of bacterial load around 1 log_10_ when compared to absence of BLIs (Fig. 6).

**Figure 6:**
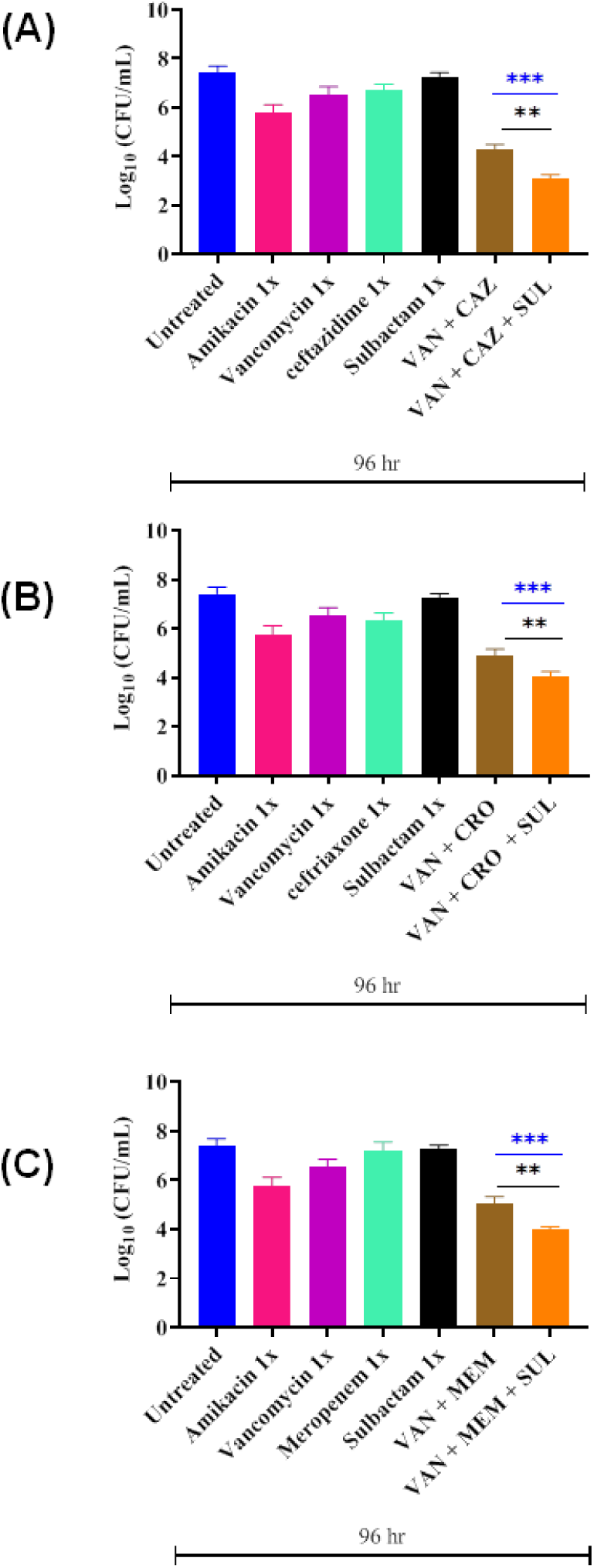
Intracellular efficacy of Vancomycin with different β-lactams in presence or absence of β-lactamase inhibitor Sulbactam (SUL) against *M. abscessus* ATCC 19977 was done in the single set of experiments for simplicity of comparison the outcomes were displayed in separate panels. Combination of VAN and BLs in presence or absence of SUL showed significantly reduced intracellular bacterial burden as compared to untreated as well as all the drugs alone; the significance value was depicted in blue stars (***, P < 0.001). Addition of SUL led to further decrease of bacterial load around 1 log_10_ CFU/mL when compared to absence of BLIs depicted in black stars (**, P value < 0.01). All values are depicted as the mean ± standard deviation (SD).

### *In vivo* efficacy of combinatorial approach against Mtb H37Rv ATCC 27294 and *M. abscessus* ATCC 19977

The *in vivo* efficacy was tested against Mtb H37Rv ATCC 27294 BALB/c in murine infection model (28). Briefly, mice were infected through IV via tail vein, kept for 7 days to allow the infection to establish and then treated with drugs alone and combinations and results are plotted in Fig. 7 A-C. As can be seen, treatment with VAN (300 mg/kg) caused a reduction of ∼0.9 log_10_ CFU/mL in lung, in kidney 0.8 log_10_ CFU/mL and in spleen 0.9 log_10_ CFU/mL compared to untreated. Whereas, both RIF (10 mg/kg) and INH (10 mg/kg) reduced bacterial load by ∼2. log_10_ CFU/mL in lung, 2.6 log_10_ CFU/mL in kidney and 2.9 log_10_ CFU/mL in spleen. EMB (100 mg/kg) reduced bacterial load by ∼2.2 log_10_ CFU/mL in lung, 0.7 log_10_ CFU/mL in kidney and 2 log_10_ CFU/mL in spleen. Treatment with various BLs and SUL alone caused a reduction of ∼0.95-1.11 log_10_ CFU/mL in lung, 0.7-1.1 log_10_ CFU/mL in kidney, 0.87-1.1 log_10_ CFU/mL in spleen as compared to untreated at 35 dpi. In comparison, treatment with VAN + MEM, VAN + CRO and VAN + CAZ caused a significant reduction ∼3.17-4.17 log_10_ CFU/mL, ∼3.1-3.6 log_10_ CFU/mL, ∼3.88-4.48 log_10_ CFU/mL as compared to untreated in lung, kidney and spleen respectively. However, there was no significant difference after addition of SUL to the combinations of VAN + BLs in any of the organs compared to the absence of SUL. Most importantly, treatment with VAN + BLs or VAN + BLs + SUL caused ∼1 to 1.5 log_10_ CFU/mL reduction in bacterial load compared to first-line anti TB drugs, INH and RIF.

**Figure 7:**
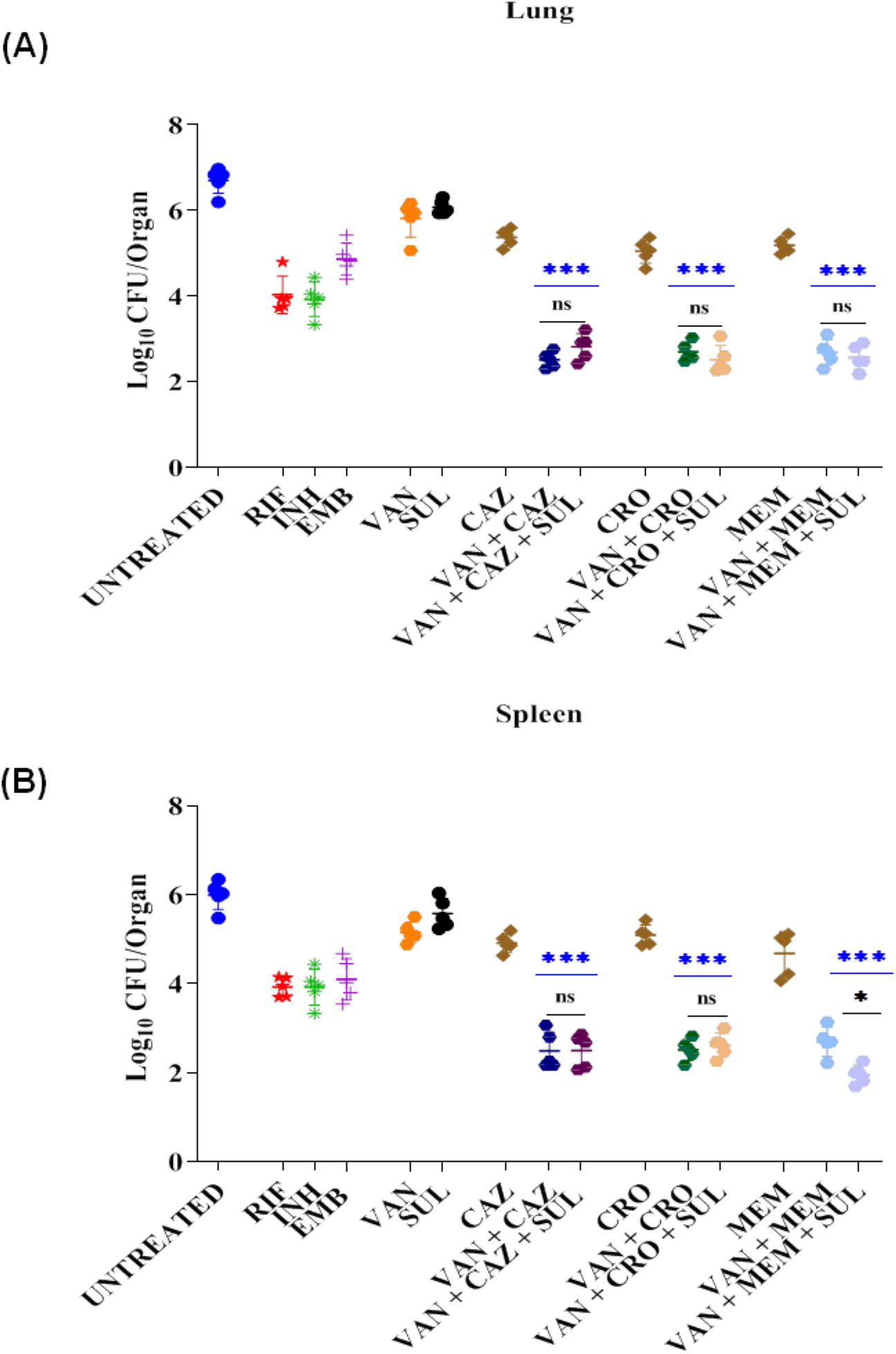
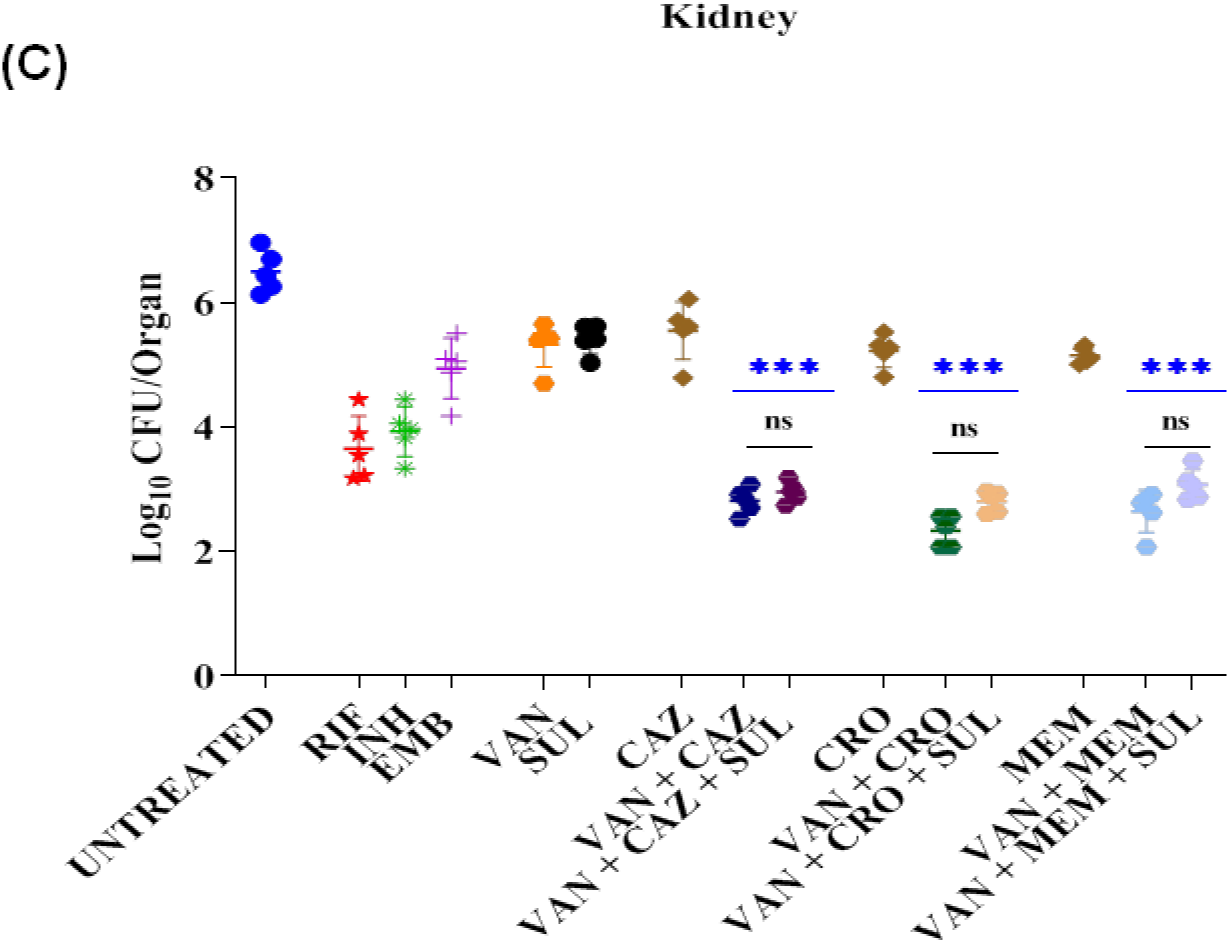
*In vivo* efficacy of Vancomycin and β-lactams alone and in combinations against Mtb H37Rv ATCC 27294. Bacterial burden different organs in the murine bacteremia model. The male BALB/c mice were infected with 5×10^5^ CFU/mL inoculum and treated with RIF (10mg/Kg), INH (10mg/Kg) & EMB (100mg/Kg) which are positive control, Vancomycin (300mg/Kg), β-lactams (100mg/Kg) and SUL (50mg/Kg) for 14 days. The bacterial burden in various organs infected with a Mtb H37Rv ATCC 27294 (∼5 × 10^5^ CFU/mL) decreased significantly after treatment with VAN + β-lactams or VAN + β-lactams + SUL, as compared to untreated drug alone or positive control; RIF, INF & EMB; significance was shown in blue stars (***, P < 0.001). In lung & kidney, combinations of VAN + BLs in presence of SUL doesn’t any significant reduction (ns, non significant) but in spleen, presence of SUL with VAN + MEM shows significant; shown in black star (*, P < 0.01) log_10_ CFU/mL decrease in comparison to VAN + MEM. All data are depicted as the mean ± SD.

Similarly, *in vivo* efficacy was also tested in the murine neutropenic *M. abscessus* ATCC19977 bacteraemia model as it mimics clinical representation of NTM infection and results are shown in Fig.8. A-C. As can be seen, treatment with VAN (300 mg/kg) and AMK reduced bacterial load by ∼1.5 log_10_ CFU/mL while treatment with BLs and SUL alone caused reduction of ∼0.74–1.4 log_10_ CFU/mL in lung at 10 dpi as compared to untreated. In comparison, the combination of VAN + BLs significantly more effective with ∼3 - 3.6 log_10_ CFU/mL reduction in lungs, kidney and spleen compared to untreated. Addition of SUL in the combinations of VAN+BLs did not show any significant improvement in the outcome compared to absence the BLI. The combinations of VAN+BLs (or VAN + BLs + SUL) comprehensively outperformed AMK with a significantly higher reduction in bacterial load (∼2 log_10_ CFU/mL) in all the organs. Collectively, the identified combinations of VAN with BLs have all the potential to be used in the for drug-resistant mycobacterial pathogens.

**Figure 8:**
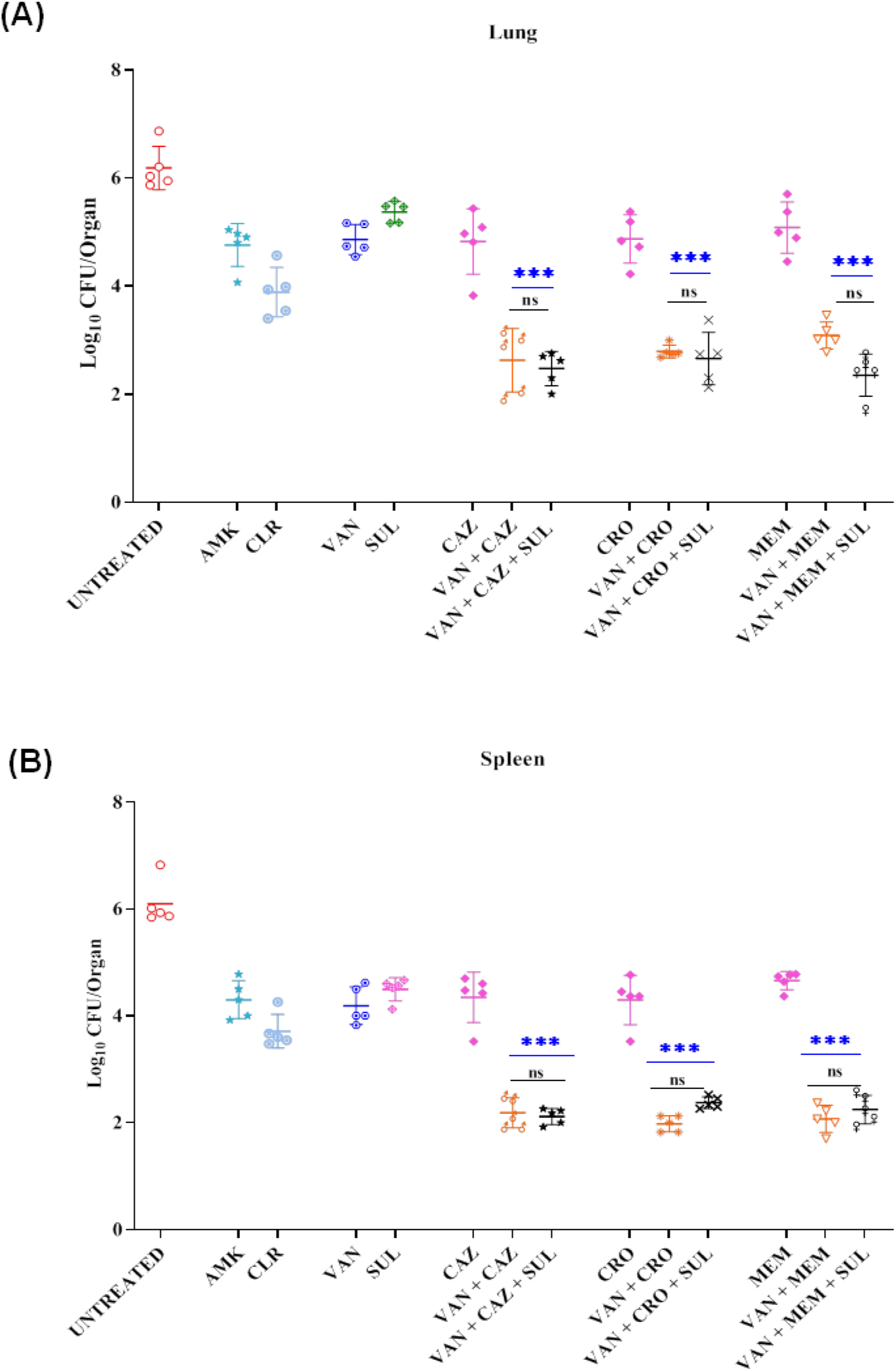
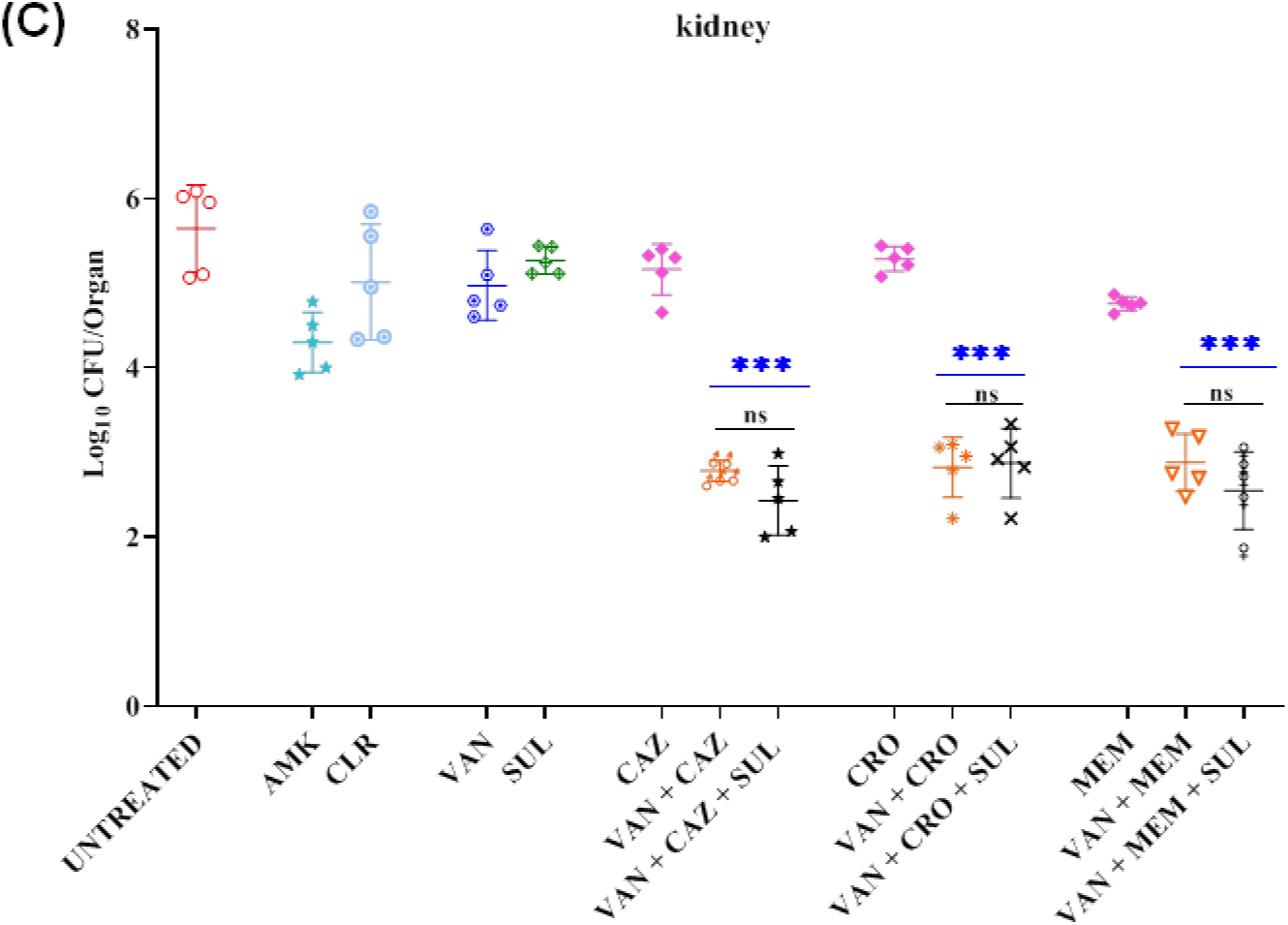
*In vivo* efficacy of Vancomycin and β-lactams alone in combinations against *M. abscessus* ATCC 19977. Bacterial burden different organs (A. Lung, B. Kidney, C. Spleen.) in the murine bacteremia model. The male BALB/c mice were infected with 5×10^5^ CFU/mL inoculum and treated with AMK (100 mg/kg) which is as positive control, Vancomycin (300mg/Kg), β-lactams (100mg/Kg) and SUL (50mg/Kg) for 10 days. After treatment the bacterial burden in various organs infected with a *M. abscessus* ATCC 19977 decreased significantly after treatment with VAN + β-lactams or VAN + β-lactams + SUL, as compared to untreated and drug alone (VAN, CAZ, CRO and SUL) or positive control AMK; significance was shown in blue stars (***, P < 0.001). In lung, kidney & spleen, combinations of VAN + BLs in presence of SUL doesn’t any significant reduction (ns, non significant). All data are depicted as the mean ± SD.

## Discussion

Since the discovery of penicillin, inhibition of PG biosynthesis has been a very attractive, highly conserved and broad-spectrum target with multiple drugs in clinical utilization inhibiting the pathway (29). In spite of their extensive clinical application, glycopeptides and β-lactams has not been extensively utilized for treatment of TB and infections caused due to NTM, mainly because of intrinsic resistance linked to low permeability, which is due to nature of complex mycobacterial cell wall (30,30). In this vein, some of the recent publications including Shin et al showed the efficacy of different BLs and BLIs against mycobacterial pathogens (22). However, in most of these works only *in vitro* activity of the combinations were shown. Herein, we offer *in vivo* proof that the drug combinations offer excellent potential for treatment of drug-resistant mycobacterial infections by increasing the permeability of mycobacterial cell, thus negating a significant resistance mechanism. Indeed, experiments carried out in *M. smegmatis*, slow growing BCG and Mtb have demonstrated that changes in cell permeability drive susceptibility to β-lactams (32–34). This low permeability is also a significant resistance mechanism in NTM’s, especially in case of macrolides and aminoglycosides (35,36,).

VAN has been used efficaciously against various gram-positive pathogens but is ineffective against gram-negative bacteria. This is primarily due to their lipid rich, outer membrane which does not permit this large hydrophilic molecule to enter (14–15). Similarly, mycobacteria are not considered susceptible to VAN, despite reports that mention successful clinical utilization, mostly against NTMs (37–39). VAN binds to D-alanine–D-alanine terminal amino acids of peptide side chains of PG and prevents the cross-linking by PBP D, L-transpeptidase and other processes in final assembly of mature PG (40). Similar to β-lactams, cell wall permeability is the primary resistance mechanism to VAN. In this context, lipid-lowering agents increase VAN susceptibility in mycobacteria by modifying permeability (41). Indeed, MIC of VAN against mycobacteria can be reduced in combination with cerulenin, again by modifying permeability (42). Further, efficacy of VAN has been assessed in alternate delivery methods including inhalable delivery against mycobacterial diseases as well as conjugating with arginine to enhance entry into both Gram positive and Gram negative bacteria including mycobacteria (42–44).

β-lactams inhibit PBPs and L,D-transpeptidases, which are involved in peptidoglycan biosynthesis thus promote the disruption of peptidoglycan layer of bacterial cell wall. Mtb produces seven different types of PBPs that include two class A, two class B and three class C. In contrast, *Mycobacterium abscessus* possesses two class A PBPs (PonA2 and PonA1) and one class B PBP (PbpA) (23). In addition to inhibiting β-lactamase activity, Sulbactam binds to PBPs, thus enhancing bacterial killing. Here, we demonstrate that combination of VAN with BLs in presence or absence of Sulbactam increase permeability of mycobacterial cell wall and show synergistic activity against drug-resistant mycobacterial pathogens. However, addition of Sulbactam did not enhance the efficacy of the synergistic combination *in vivo* compared to VAN and BLs.

One of the most striking advantages of this combination is that it exhibits high efficiency in killing combined with low rate of emergence of mutants. The clinical strains of mycobacteria exhibit little variation in β-lactamase activity, there is no evidence of VanA and VanB mediated resistance against VAN and chances of acquired resistance are low because mycobacteria have very low potential of horizontal gene transfer (40–42). Thus, it seems fairly possible that resistance to this combination can be thwarted for a reasonable period of time. Dosing regimens would need to be determined to ensure optimal drug exposures.

## Conclusion

This proof-of-concept study identifies a highly efficacious synergistic combination of glycopeptides, BLs and a BLI to combat drug resistant mycobacterial pathogens including *M. tuberculosis* and notoriously MDR *M. abscessus*. The combinations demonstrate significant potential to control mycobacterial growth both *in vitro* and *in vivo* that comprehensively outperformed first-line clinically utilized antibiotics such as INH, RIF and EMB against *M. tuberculosis* and AMK. However, addition of BLI in the combination of VAN and BLs demonstrated significantly higher efficacy, but it did not translate to increased *in vivo* efficacy. A detailed systematic study is underway to identify the best combinations of VAN, BL and BLI for optimized clinical application against MDR mycobacterial infections.

## Material and Methods

### Chemicals and Reagents

All bacterial media and supplements including Middlebrook 7H9 broth, Middlebrook 7H11 agar, ADC and OADC (Oleic acid, Albumin, Dextrose and Catalase) supplements were purchased from BD (Franklin Lakes, NJ, USA). All the other chemicals and antibiotics were procured from Sigma-Aldrich (St. Louis, MO, USA).

### Bacterial cultures and animal cell line

The following mycobacterial strains were meticulously obtained from the American Type Culture Collection (ATCC, Manassas, USA): drug-susceptible Mtb H37Rv (ATCC 27294), Isoniazid (INH)-resistant Mtb (ATCC 35822), Rifampicin (RIF)-resistant Mtb (ATCC 35838), Streptomycin (STR)-resistant Mtb (ATCC 35820), Ethambutol (EMB)-resistant Mtb (ATCC 35837), *M. chelonae* (ATCC 35752), *M. abscessus* (ATCC 19977), *M. abscessus* (MC1815) and *M. abscessus* (DJO 4274). The mycobacterial strains were propagated in Middlebrook 7H9 broth enriched with 10% ADC, 0.2% glycerol, and either 0.1% Tyloxapol or Tween-80 at 37°C. For accurate colony- sforming unit (CFU) enumeration, we employed Middlebrook 7H11 agar supplemented with 0.2% glycerol and 10% OADC, ensuring precise results for all mycobacterial strains.

### Antibacterial susceptibility testing

Antibacterial susceptibility testing was performed utilizing broth microdilution assay according to CLSI guidelines as described previously (46). Briefly, 100 mg/mL stock solutions of test and control antibiotics were prepared in DMSO or water depends upon their solubility. Bacterial cultures were inoculated in Middlebrook 7H9 broth supplemented with 10% OADC, 0.2% glycerol, 0.05% tween-80 and OD_600_ of cultures was measured, followed by dilution to achieve ∼10^6^ CFU/mL. The compounds were tested from 1024 to 0.03 μg/mL in a two-fold serial diluted fashion with 2.5 μL of each concentration added per well of a 96-well round bottom microtiter plate. Later, 97.5 μL of bacterial suspension was added to each well containing the test compound along with appropriate controls and incubated at 37°C for 48 h for rapid-growing mycobacteria and 7 days for slow-growing mycobacteria. The MIC is defined as the lowest compound concentration with no visible growth. For each compound, MIC determinations were carried out independently 3 times in duplicate and mean value is presented.

### Synergy between Vancomycin and β-lactams

The interaction between VAN and BLs such as CRO, CAZ and MEM was tested by checkerboard method as per CLSI guidelines as described previously (46). Briefly, serial two-fold dilutions of each drug were freshly prepared before testing. VAN was two-fold diluted along the ordinate while CRO, CAZ and MEM were serially diluted along the abscissa in 96 well microtiter plate. 95 µL of ∼10^6^ CFU/mL of various mycobacterial pathogens were added to each well and plates were incubated at 37^0^C for 48 h for rapid growers and 7 days for slow growing mycobacteria. After the incubation period was over, the ΣFICs (fractional inhibitory concentrations) were calculated as follows:

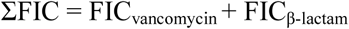

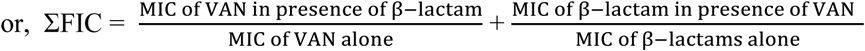

The combination is considered synergistic when the ∑FIC is ≤0.5, indifferent when the ∑FIC is >0.5 to 4, and antagonistic when the ∑FIC is >4 (47).

### Combinatorial time kill assay

To determine whether the combination of VAN with BLs in presence or absence of BLI exhibit bactericidal or bacteriostatic activity time-kill assay was performed (46, 48,49). The mycobacterial cells were diluted to achieve ∼10^6^ CFU/mL and added to 96 well plate along with various antibiotics and controls at 1x MIC, followed by incubation at 37^0^C for 72 h for rapid growing mycobacteria and 7 days for slow growing mycobacteria. For evaluating the reduction in CFU, 30 µL sample was removed at various time points, serially two-fold diluted in 270 µL normal saline and 100 µL of the respective dilution was spread on Middlebrook 7H11 agar plate supplemented with 0.2% glycerol and 10% OADC. The plates were incubated at 37^0^C for 72 h for rapid growing mycobacteria and 28 days for slow growing mycobacteria, after which colonies were enumerated. Kill curves were constructed by counting the colonies from plates and plotting the CFU/mL of surviving bacteria at each time point in the presence and absence of the compound. Each experiment was repeated three times in duplicate and the mean data is plotted.

### Ethidium bromide (EtBr) assay to study cell wall permeability

*M. abscessus* ATCC 19977 were grown to an OD_600nm_ of 0.4 - 0.6 in 7H9 media supplemented with ADC and 0.05% Tween-80 and used for an EtBr uptake assay as described elsewhere (40,50). Briefly, *M. abscessus* cultures were dispensed in opaque-walled, 96-well, flat-bottomed cell culture plates and pre-energized with 0.4% glucose for 5 min. Thereafter, drugs alone and in combination (at ½ MIC to not compromise cellular viability) were added, followed by addition of 10 μM EtBr. The relative fluorescence units (RFUs) were monitored every two minutes by Excitation (530 nm)/ Emission (590 nm) filter using multi-mode plate reader for 60 min and the results were plotted as change in fluorescence intensity (OD).

### Intracellular killing assay against *Mycobacterium abscessus*

The efficacy of the combinations against intracellular *M. abscessus* was performed as described earlier (46,48). Briefly, *M. abscessus* ATCC 19977 was culture in supplemented 7H9 Middlebrook medium for overnight to rich mid-log phase. From the same culture bacterial inoculum was prepared in 1x phosphate saline buffer (PBS) with ∼ 10^7^ CFU/ml. Macrophage cell line J774A.1 were seeded in 12 well flat bottom plates at a density of ∼ 10^5^ cells/well and after 24 hours macrophage cells were infected with *M. abscessus* at a 1:5 multiplicity of infection (MOI). After 4 hours of post-infection the cells were washed twice with 1x PBS and resuspended with fresh RPMI medium containing either VAN, BLs and sulbactam alone, Vancomycin combination with beta-lactams or vancomycin combination with beta-lactams in presence of sulbactam and Amikacin use as a control. To enumerate the initial bacterial load three wells were lysed after 4 hours of post-infection, and lysates were serially diluted and plated on supplemented 7H11 agar plates. After 96 hours of incubation, another set of 3 wells of each group were lysed, serially diluted, plated on 7H11-ADC agar plates and incubated at 37°C for 72 h to estimate the CFU. The reduction of the bacterial load was plotted in CFU/mL by counting colonies from the agar plate. Each experiment was carried out three times in replicate, and the mean value was plotted.

### Enumeration of live/dead bacteria using fluorescent microscopy

The viability of *M. abscessus* ATCC 19977 after treatment with VAN, MEM, and SUL alone or in combination with VAN+ MEM or VAN + MEM + SUL were evaluated using LIVE/DEAD BacLight bacterial viability kit (Invitrogen, catalogue No. L7007) according to the manufacturer’s and published protocol (51). Briefly, *M. abscessus* ATCC 19977 cells were cultured overnight in 7H9-ADC Middlebrook broth till OD_600nm_ ∼ 0.6. The mid-log bacterial culture was then diluted to approximately ∼ 10^7^ CFU/mL in 7H9-ADC Middlebrook broth and treated with VAN, MEM, and SUL alone or VAN+ MEM or VAN + MEM + SUL at 1× MIC for 48 h at 37°C. After incubation, cells were washed two times and resuspended in Tween80 normal saline (TNS). 3 μL of propidium iodide (PI) and SYTO9 (ratio 1:1) provided in the kit were added to each sample, with a final volume of 1 mL and samples were kept at room temperature in dark condition for 15 min. fluorescent images were taken from stained samples using fluorescence microscope with 100X oil immersion objective.

### Scanning Electron Microscopy

Using Scanning Electron Microscopy (SEM) the effect of combinations on topological and morphological alterations in *M. abscessus* was observed. The mid-log phase (OD_600nm_ = 0.6-0.7) *M. abscessus* (ATCC19977) culture was treated with 5x MIC of VAN and BLs alone and VAN + BL combinations for 8 h at 37°C. After incubation treated cells were washed three times with 1x PBS and proceeded as described previously (46, 52). The washed cells were fixed with 2.5% glutaraldehyde made in 0.1 M phosphate buffer. The cell suspensions were placed on poly-l-lysine-coated glass chips after washing with the same PBS and kept for 10 min at room temperature to attach to the glass chips. Hereafter, the samples were postfixed in 1% Osmium tetroxide (OsO4) and consequently dehydrated through an increasing concentration of ethanol series, were critical point dried. Then the samples were coated with a mixture of Au-Pd (80:20) using a Polaron E5000 sputter coater and examined under FEI Quanta 250 scanning electron microscope (SEM) at an accelerating voltage of 10 kV, using the Secondary detector.

### Animal experiments

Animal experiments were performed on six-eight-week-old BALB/c female mice procured from the National Laboratory Animal Facility of CSIR-Central Drug Research Institute, Lucknow. The experimental protocols were approved by the Institutional Animal Ethics Committee. Animal experiments were performed per the guidelines provided by the Committee for Control and Supervision of Experiments on Animals (CPCSEA, Govt. of India).

### *In vivo* determination of efficacy of combination therapy against Mtb H37Rv ATCC 27294

6-8-week-old, female, BALB/c mice were infected with ∼5×10^5^ CFU/mL of Mtb H37Rv ATCC27294 through intravenously (IV) in the tail vein and divided into 15 groups randomly, with each group having 6 mice and the untreated group receiving 12 mice as described previously (28). After seven days of post-infection (dpi), drug treatment was initiated according to Table 3A:

| Groups | Treatment | Dosage (mg/kg) |
| --- | --- | --- |
| 1. | Untreated (PBS) | NA |
| 2. | RIF | 10 |
| 3. | INH | 10 |
| 4. | EMB | 100 |
| 5. | VAN | 300 |
| 6. | SUL | 50 |
| 7. | CAZ | 100 |
| 8. | VAN+CAZ | 300+100 |
| 9. | VAN+CAZ+SUL | 300+100+50 |
| 10. | CRO | 100 |
| 11. | VAN+CRO | 300+100 |
| 12. | VAN+CRO+SUL | 300+100+50 |
| 13. | MEM | 100 |
| 14. | VAN+MEM | 300+100 |
| 15. | VAN+MEM+SUL | 300+100+50 |

All drugs were prepared in PBS. INH, RIF, and EMB were administered 200 µl orally while all other drugs were administered 100 µl intraperitoneally (IP). 6 mice were sacrificed at 7 dpi before treatment to evaluate the early load of infection. After 4 weeks post-treatment, mice were sacrificed, spleens, kidneys and lungs were recovered, homogenized in 5 ml of PBS, serially diluted and plated onto Middlebrook 7H11 and the CFU of each mouse was plotted.

### *In vivo* determination of efficacy of combination therapy against *M. abscessus* ATCC 19977

6-8-week-old, female, BALB/c mice were infected IV with ∼5×10^5^ CFU/mL of *M. abscessus* ATCC 19977 and were divided into thirteen groups randomly, with each group having 6 mice and the untreated group receiving 12 mice as described previously (4). After 3 dpi, drug treatment was initiated according to Table 3B:

| Groups | Treatment | Dosage (mg/kg) |
| --- | --- | --- |
| 1. | Untreated (PBS) | NA |
| 2. | AMK | 100 |
| 3. | VAN | 300 |
| 4. | SUL | 50 |
| 5. | CAZ | 100 |
| 6. | VAN+CAZ | 300+100 |
| 7. | VAN+CAZ+SUL | 300+100+50 |
| 8. | CRO | 100 |
| 9. | VAN+CRO | 300+100 |
| 10. | VAN+CRO+SUL | 300+100+50 |
| 11. | MEM | 100 |
| 12. | VAN+MEM | 300+100 |
| 13. | VAN+MEM+SUL | 300+100+50 |

All drugs were prepared in PBS, administered 100 µl intraperitoneally (IP) and 12 untreated mice received PBS only. All drugs were given single dose/day till 10 days of post treatment. Six mice were sacrificed at 3 dpi to enumerate the initial load. After 10 days post treatment, mice were sacrificed, spleens, kidneys, and lungs were recovered, homogenized in 5 ml of PBS, serially diluted and plated onto Middlebrook 7H11 for enumeration of CFU.

### Statistical analysis

Statistical analysis was performed using GraphPad Prism software (GraphPad Software, La Jolla, CA, USA). Comparison between three or more groups was analyzed using one-way ANOVA, with post-hoc Tukey’s multiple comparisons test. P-values of <0.05 were considered to be significant.

## Acknowledgement

SC and AD thanks ICMR (ICMRCAREP-2023-0000461) for funding. AKS, MI and MNA thank CSIR while PM, AM and JA thank UGC for their fellowships. The authors thank Mr. Jeevan Prakash Pandey for his technical help with scanning electron microscopy. This manuscript bears CSIR-CDRI communication number XXXX.

## Authors’ contributions

AKS, PM, SC and AD conceptualized the work, AKS, PM, UDG, KM, SC and AD designed experiments, AKS, PM, AM, MI, MNA, JA, RM, SP, AG and PK performed experiments; AKS, PM, UDG, KM, SCand AD analyzed data; AKS, PM, UDG, KM, SC and AD wrote and edited the manuscript. All authors reviewed and approved the manuscript.

